# Targeting tumor-stromal interactions in triple-negative breast cancer using a human vascularized micro-tumor model

**DOI:** 10.1101/2023.09.06.556584

**Authors:** Stephanie J. Hachey, Christopher J. Hatch, Daniela Gaebler, Aneela Mocherla, Kevin Nee, Kai Kessenbrock, Christopher C.W. Hughes

## Abstract

Triple-negative breast cancer (TNBC) is highly aggressive with limited available treatments. Stromal cells in the tumor microenvironment (TME) are crucial in TNBC progression; however, understanding the molecular basis of stromal cell activation and tumor-stromal crosstalk in TNBC is limited. To investigate therapeutic targets in the TNBC stromal niche, we used an advanced human *in vitro* microphysiological system called the vascularized micro-tumor (VMT). Using single-cell RNA sequencing (scRNA-seq), we revealed that normal breast-tissue stromal cells activate neoplastic signaling pathways in the TNBC TME. By comparing interactions in VMTs with clinical data, we identified therapeutic targets at the tumor-stromal interface with potential clinical significance. Combining treatments targeting Tie2 signaling with paclitaxel resulted in vessel normalization and increased efficacy of paclitaxel in the TNBC VMT. Dual inhibition of Her3 and Akt also demonstrated efficacy against TNBC. These data demonstrate the potential of inducing a favorable TME as a targeted therapeutic approach in TNBC.

## 1 Introduction

Triple-negative breast cancer (TNBC), representing approximately 15% of all breast carcinomas, is an aggressive disease with limited treatment options [1]. Consequently, TNBC disproportionally accounts for 30% of all breast cancer-related mortality, with approximately 150,000 deaths estimated in 2023 [2]. Since TNBC, by definition, lacks expression of the estrogen receptor (ER), progesterone receptor (PR), and human epidermal growth factor receptor 2 (HER2), chemotherapy remains standard for systemic treatment, as endocrine therapy and HER2-directed therapies are ineffective [3]. Compared with hormone receptor (HR)-positive breast cancer, TNBC is more commonly diagnosed in women younger than 40 years and has a worse 5-year survival rate [3]. A study evaluating 15,204 women who presented with stage I-III breast cancer at National Comprehensive Cancer Network Centers found that women with TNBC experienced worse breast cancer-specific and overall survival (OS) in multivariate analyses (OS: adjusted hazard ratio, 2.72; 95% confidence interval, 2.39-3.10; P *<*0.0001) compared to women with HR-positive/HER2-negative breast cancer [4]. Additionally, TNBC tumors have been found to be associated with unique risk factors and poor prognosis, including an increased risk of death within 2 years of diagnosis [4], highlighting the need to develop more effective treatments for this deadly malignancy.

Current first-line therapy for patients with TNBC includes neo/adjuvant anthracycline-, alkylator-, and taxane-based chemotherapy regimens, for example doxorubicin and cyclophosphamide followed by paclitaxel [5]. However, chemotherapy achieves a complete response in only about 30% of TNBC patients, and more than 50% of patients experience relapse within the first 3-5 years due to the development of therapy resistance. Additionally, patients with stage IV TNBC treated with standard-of-care chemotherapy have a median overall survival of just 10.2 months [5]. There is growing evidence that stromal cells in the tumor microenvironment (TME) play an important role in cancer progression and treatment resistance in TNBC [5–9], and are therefore considered a potential therapeutic target. The TME, a major determinant of cancer phenotype, is a complex and heterogeneous ecosystem co-opted by neoplastic cells, consisting of both cancerous and nonmalignant cell populations embedded in a glycoprotein-rich extracellular matrix (ECM). Cancer-associated fibroblasts (CAFs) are a prominent component of the tumor stroma from which gene expression signatures have been found to strongly predict TNBC prognosis. Upregulated pathways include those associated with ECM remodeling and chemokine signaling, which contribute to tumor cell growth and invasion as well as vascular destabilization [10–12]. However, the underlying molecular mechanisms driving stromal cell activation and the reciprocal communication between tumor cells and stromal cells in TNBC remain poorly understood, impeding the development of targeted therapies focused on the stromal component.

In this study, our aim was to uncover mechanisms by which the microenvironment of TNBC triggers the reprogramming of stromal cells into an activated state, ultimately influencing both tumor growth and response to therapy. Additionally, we aimed to identify and evaluate potential therapeutic targets for TNBC. Herein we establish human vascularized micro-tumor (VMT) models to recreate the TNBC TME *in vitro*, using primary stromal cells isolated freshly from healthy human breast tissue, endothelial cells (ECs) and two representative TNBC cell lines, HCC1599 and MDA-MB-231. The VMT is a microphysiologic *in vitro* cancer model that we have validated for disease modeling, drug screening and personalized medicine applications for solid tumors [13–17]. By co-culturing multiple cell types within a complex ECM under dynamic flow conditions, the stromal cells and endothelium spontaneously self-organize into a perfused vascular network that plays a vital role in supporting tumor growth and serves as a physiological conduit for the delivery of therapeutic agents to the tumor. Furthermore, the VMT accurately replicates key characteristics of an *in vivo* tumor, including the presence of irregular and poorly perfused tumor-associated vasculature, which can hinder the effective delivery of therapeutics to the tumor and contribute to treatment resistance [16].

By leveraging single-cell RNA sequencing (scRNA-seq), we investigated gene expression patterns, cellular heterogeneity, and intercellular communication within TNBCs. Our analysis revealed that histopathologically normal human breast-tissue derived stromal cells activate neoplastic signaling pathways when exposed to the TNBC TME. Through a comparison of these interactions in VMTs with scRNA-seq data obtained from clinical specimens, we successfully identified therapeutic targets situated at the tumor-stromal interface that hold potential clinical significance.

## 2 Results

### Healthy breast tissue-derived stromal cells support microvascular network formation in VMO and VMT models

For physiologic modeling of TNBC, primary-derived stromal cells were freshly isolated from histopathologically normal (healthy, noncancerous) breast tissue under an IRB-approved protocol and tested for vascular network formation in VMO/VMTs (Figure 1). VMOs and VMTs are established within a microfluidic device (Figure 1A) via coculture of multiple cell types within an extracellular matrix. Cells are introduced into the tissue chamber of each unit through the loading port (L1 or L2) and experience dynamic flow via a hydrostatic pressure gradient through the microfluidic channels from the media reservoirs (M1-M2 and M3-M4). In response to flow across the tissue chamber, ECs and stromal cells self-assemble into a microvascular network by day 5 of culture to allow *in vivo*-like delivery of nutrients and drugs to the tissue.

**Fig. 1:**
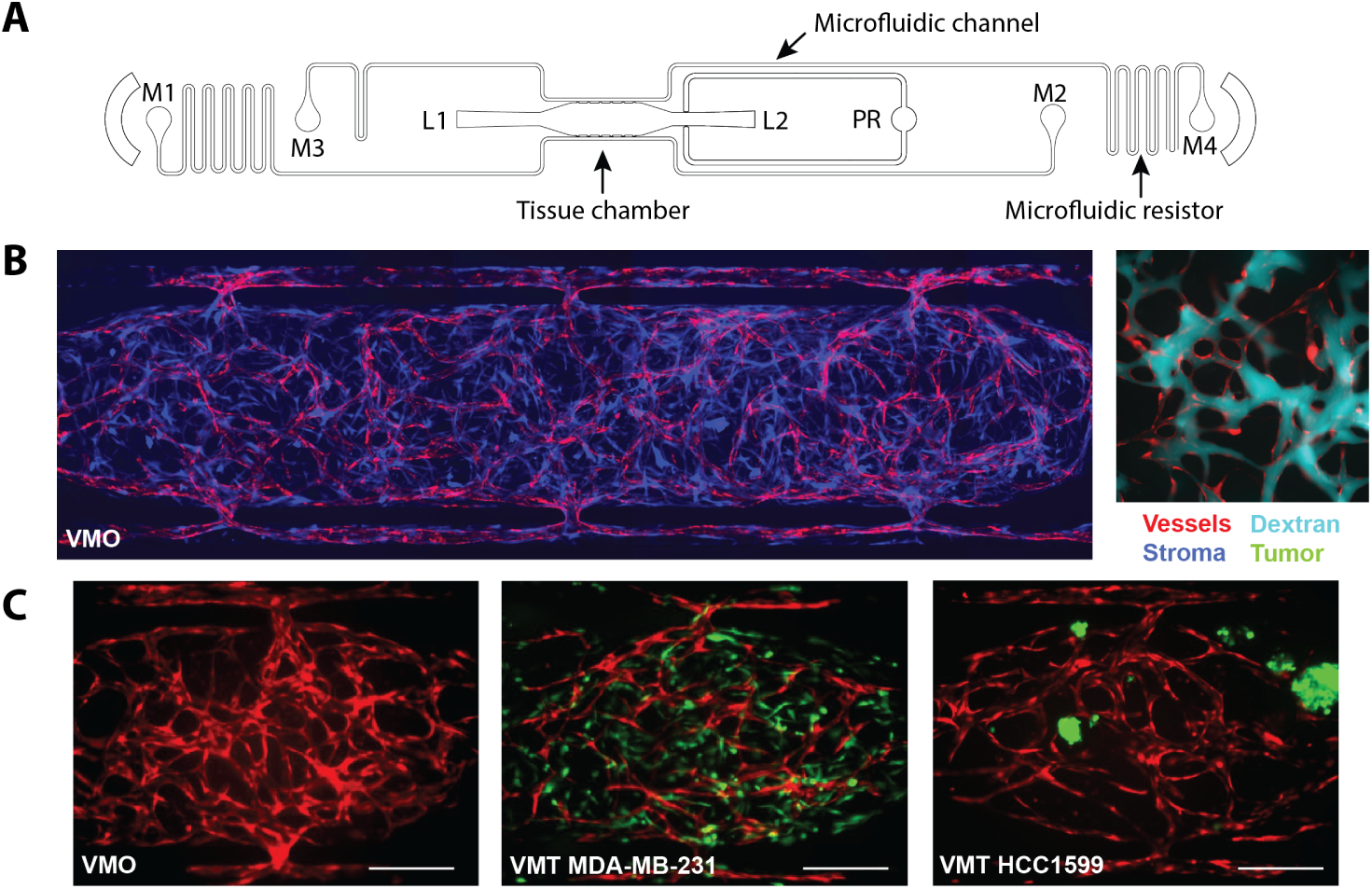
Stromal cells freshly isolated from healthy breast tissue support microvascular network formation in VMO and VMT models. **(A)** Schematic of a single device unit with a single tissue chamber fed through microfluidic channels, 2 loading ports (L1-2), and uncoupled medium inlet and outlets (M1-2 and M3-4). A pressure regulator (PR) serves as a burst valve to release excess pressure from the tissue chamber during loading. **(B)** Left: Fluorescent image of a VMO established with stromal cells isolated freshly from histopathologically normal breast tissue. Vessels are shown in red and primary fibroblasts in blue. Right: Zoom view of a vascular network (red) perfused with 70 kDa dextran (cyan). **(C)** Left panel: VMO established with patient-derived breast-tissue stromal cells. Vasculature shown in red. Middle panel: VMT established with MDA-MB-231 TNBC cells and patient-derived stromal cells. Note invasiveness of the tumor cells. Vasculature is red, tumor is green, and stromal cells are unlabeled. Right panel: VMT established with HCC1599 TNBC cells and patient-derived stromal cells. Vasculature is red, tumor is green, and stromal cells are unlabeled. Scale bar = 200 µm.

As shown in Figure 1B, healthy breast tissue-specific primary stromal cells (shown in blue) support the development of microvessels (red) that are fully perfused by 70 kDa fluorescent dextran (cyan). Matched VMOs and VMTs containing TNBC cell lines MDA-MB-231 and HCC1599 (shown in green) were established (Figure 1C). MDA-MB-231 grows rapidly with stellate morphology and manifests invasive spread within the VMT, gradually disrupting the surrounding vasculature over time. In contrast, HCC1599 exhibits slow proliferation as 3D clusters and has been demonstrated to closely resemble clinical TNBC samples in terms of various molecular characteristics, such as gene expression signatures and copy number variations [18, 19].

### Single-cell RNA sequencing reveals differential pathway activation in TNBC VMTs and VMOs

In order to gain deeper insights into the alterations of the vascular niche in TNBC tumors, we conducted scRNA-seq on cells isolated from TNBC VMTs (MDA-MB-231 and HCC1599) and their corresponding VMOs (Vasculature and stroma in the absence of tumor cells; Figure 2A). Using the SCTransform Seurat pipeline [20], we performed clustering analysis and confirmed distinct clusters corresponding to ECs, various stromal cells, and, in the case of TNBC VMTs, tumor cells for each dataset (Figure 2B-2I). The identification of cell types was determined based on top differentially expressed genes and the expression of known marker genes specific to each cell type. These included, but were not limited to: *PECAM1* and *CDH5* (EC markers), *PDGFRA* (fibroblast marker), *PDGFRB* (pericyte marker), and *EPCAM* and *KRT18* (cancer cell markers). Additionally, the matched MDA-MB-231 VMT and VMO datasets exhibited a population of cycling cells characterized by increased expression of *TOP2A* (Figure 2B-2E). Although there are slight differences in the UMAP and clustering results between the two VMOs (each VMO established at the same time as its matched VMT – MDA-MB-231 or HCC1599 VMTs), likely due to batch effects, integration of the datasets using scMC [21] reveals a seamless overlap between the cell populations (supplemental figure A1A). Both VMOs show representation of ECs, fibroblasts, and stromal cells (supplemental figure A1B), demonstrating that there were no significant differences between the VMOs.

**Fig. 2:**
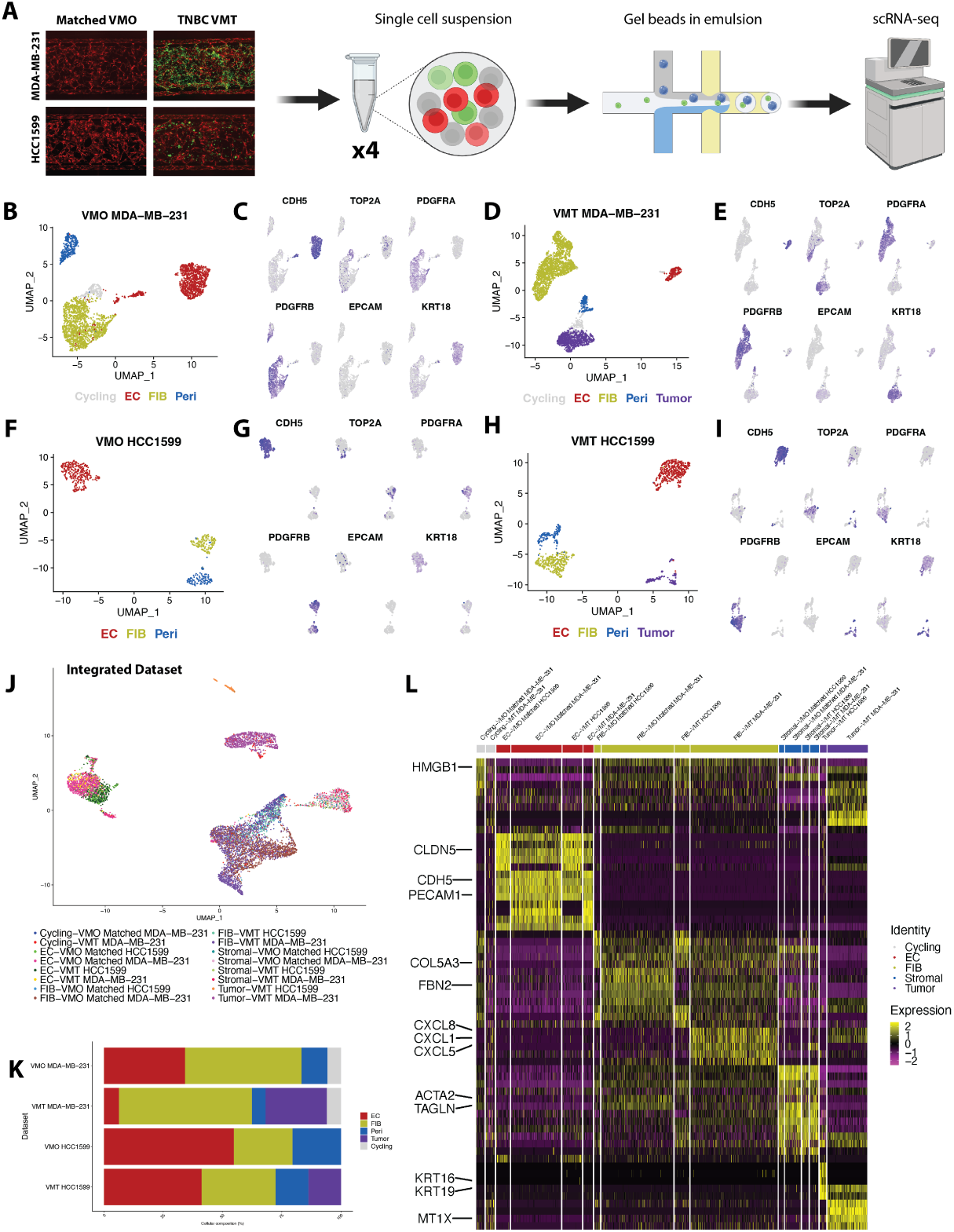
Single-cell RNA sequencing reveals transcriptomic shifts in triple negative breast cancer VMTs and VMOs. **(A)** Schematic showing workflow of scRNA-seq. Matched VMO and VMT were established for TNBC cell lines MDA-MB-231 and HCC1599. Cells were isolated from each model and submitted for scRNA-seq. **(B)** UMAP of VMO matched to MDA-MB-231 VMT shows endothelial cells (ECs), fibroblasts, cycling cells and pericytes are present as **(C)** shown with the expression of key marker genes. **(D)** UMAP of MDA-MB-231 VMT shows ECs, fibroblasts, pericytes, cycling cells and tumors are present as **(E)** demonstrated with the expression of key marker genes. **(F)** UMAP of VMO matched to HCC1599 shows ECs, fibroblasts, and pericytes are present as **(G)** demonstrated with the expression of key marker genes. **(H)** UMAP of HCC1599 VMT shows ECs, fibroblasts, pericytes, and tumors are present as **(I)** shown with the expression of key marker genes. **(J)** Integrated dataset containing VMO (MDA-MB-231), VMO (HCC1599), MDA-MB-231 VMT and HCC1599 VMT. **(K)** Proportions of different cell types from each dataset. **(L)** Heatmap showing top 5 DEGs from each cell type in the integrated dataset.

Next, to examine the variations between datasets, all four datasets were integrated using scMC [21]. Unsupervised clustering of the integrated data revealed distinct clusters corresponding to the cell identities defined in the individual datasets, with clear separation of tumor cell lines (Figure 2J). When analyzing the proportion of cell types contributed by each sample, we observed an increase in the percentage of fibroblasts and cancer cells in MDA-MB-231 VMTs compared to HCC1599 VMTs, accompanied by a decrease in the number of ECs (Figure 2K). Comparing VMTs with VMO controls, HCC1599 VMT exhibited a marginal decrease in the number of endothelial cells, while MDA-MB-231 VMT displayed a significant decrease. These findings support the phenotypic results, where we observed a significant disruption and pruning of tumor-associated vasculature in the presence of MDA-MB-231 TNBC. That said, a heatmap depicting the top five differentially expressed genes across datasets did demonstrate consistent upregulation of *VWF* and *CLDN5* among ECs (Figure 2L). On the other hand, in contrast to HCC1599 VMT and VMOs, fibroblasts from MDA-MB-231 VMTs exhibited heightened expression of pro-inflammatory markers, including *CXCL1*, *CXCL5*, *CXCL8*, and *IL1RL1*.

### Differential gene expression analyses of individual datasets reveal distinct pathway activation dependent on cell type in both VMTs and VMOs

Both VMTs and their corresponding VMO controls demonstrate cell-type specific gene expression and pathway activation with some distinctions (supplemental figure A1C-A1F, supplemental table 1). Notably, MDA-MB-231 VMT ECs have increased cell and tissue migration and a marginal decrease of angiogenesis, most prominently noted by a lack of CXCR4, which is essential for EC tube formation [22] (supplemental figure A1G). Although both VMO and VMT fibroblasts regulate their surrounding ECM, one gene that is unique to the VMT population is MMP2, which has been shown to have increased expression in the presence of breast cancer [23](supplemental figure A1H). In addition, the VMT stromal cells express genes known to increase neutrophil activation, which may support the dual role of neutrophil activation in promoting and suppressing cancer *in vivo* [24](supplemental figure A1I). Lastly, in the MDA-MB-231 VMT, the cycling cluster is enriched for RNA splicing, for which pathway alterations have been implicated in many human cancers (supplemental figure A1J).

In the HCC1599 VMT, the ECs exhibited a stronger influence on extracellular matrix organization, driven by increased expression of *MMP1/2* (supplemental figure A1). The HCC1599 VMT fibroblasts demonstrated deficiencies in regulating angiogenesis, as evidenced by the lack of expression of SPHK1, which has been shown to be necessary for supporting endothelial tube cell formation [25] (supplemental figure A1). Additionally, the HCC1599 VMT fibroblasts exhibited increased expression of Inhibin Beta-A (INHBA), which has been implicated in epithelial-mesenchymal transition (EMT) and correlated with decreased breast cancer survival [26]. In contrast, stromal cells in the VMO control for the HCC1599 VMT exhibited better regulation of the ECM, with increased expression of *COL1A1* and *TGFBI*, both crucial for supporting lumen formation [27].

### Tumor-specific analysis highlights differences in TNBC cell populations within the VMT

To focus on TNBC cell populations within the VMT, we integrated the MDA-MB-231 VMT and HCC1599 VMT datasets and subset the tumor cells from the EPCAM+ or KRT18+ clusters in the individual datasets for further investigation. Through unbiased clustering, we discovered heterogeneity within the tumor populations in both MDA-MB-231 and HCC1599, resulting in three distinct populations in MDA-MB-231 and two populations in HCC1599 (supplemental figure A2A). Differential gene expression analysis revealed notable differences among the cellular populations within both MDA-MB-231 and HCC1599 VMT models (supplemental figure A2B). Interestingly, in MDA-MB-231 TNBC, we observed a subset of cancer cells with a transcriptional profile resembling that of stromal cells. These cells exhibited increased expression of matrix-associated genes such as collagens (*COL1A1, COL1A2, COL3A1, COL6A1, COL6A2, COL6A3*) and fibronectin (*FN1*). Correspondingly, pathway analysis revealed an enrichment of processes related to extracellular matrix organization, collagen fibril organization, and cell-substrate adhesion (supplemental figure A2C).

The cycling population of MDA-MB-231 demonstrated elevated expression of *MKI67* and *TOP2A*, along with pathway enrichment associated with cell proliferation, including nuclear division and spindle organization (supplemental figure A2D). The TNBC population of MDA-MB-231 exhibited upregulation of *AMIGO2*, which is implicated in cancer metastasis [28], and enhanced translational initiation (supplemental figure A2E). Strikingly, and in contrast to published expression data from monolayer cultures [29, 30], MDA-MB-231 cells growing in the VMT lacked expression of the epithelial marker *EPCAM*, indicating transcriptional consistency with the invasive and migratory phenotype of MDA-MB-231 in the VMT. In contrast, HCC1599-1 displayed increased expression of basal epithelial markers (cytokeratins *KRT14*, *KRT15*, *KRT16*, *KRT17*) and *EPCAM*, accompanied by enrichment for keratinization processes (supplemental figure A2F). HCC1599-2 exhibited pathway enrichment associated with positive regulation of vascular development and extracellular matrix organization (supplemental figure A2G, aligning with the tumor growth and vascular morphology patterns observed in HCC1599 VMT.

### A population of cells with a cancer-associated fibroblast (CAF) gene signature emerges from healthy breast tissue-derived stroma in MDA-MB-231 VMT

By subsetting fibroblasts from the integrated dataset, a distinct cellular population exhibiting a gene signature associated with cancer-associated fibroblasts (CAFs) emerges from the healthy breast tissue-derived stroma (supplemental figure A3A) and demonstrates enrichment within the MDA-MB-231 VMT (supplemental figure A3B). Notably, the MDA-MB-231 VMT displays an increased proportion of synthetic fibroblasts, while the HCC1599 VMT exhibits a significant enrichment of cycling fibroblasts (supplemental figure A3B). Expression profiles of differentially expressed genes included: cycling cells (*TOP2A*, *MKI67*), fibroblasts (*GATA3*, *FLRT2*), synthetic fibroblasts (vFIB) (*COL1A1*, *PCOLCE*, *SPARC*, *IGFBP7*), and cancer associated fibroblasts (CAFs) (*CCL2*, *CXCL8*, *MMP2*) (supplemental figure A3C, supplemental table 2). Differential gene expression analyses revealed distinct patterns in gene transcription among different fibroblast subtypes (supplemental figure A3D).

CAFs exhibited heightened expression of CAF-associated genes, such as *SERPINE2* and *MMP2*, which are involved in cell chemotaxis and extracellular matrix (ECM) remodeling (supplemental figure A3E). Conversely, synthetic fibroblasts displayed elevated expression of genes related to ECM organization and collagen deposition, including *COL1A1/2*, *COL3A1*, *COL4A1*, and *COL5A1/3* (supplemental figure A3F). Cycling fibroblasts exhibited increased expression of cell cycle genes and demonstrated enrichment in mitotic processes (supplemental figure A3G). Moreover, fibroblasts exhibited upregulation of *FGF2* and biological pathways associated with responses to nutrients and external stimuli (supplemental figure A3H).

### Pseudotime analysis of the EC subset unveils a tumor-associated EC signature enrichment within the MDA-MB-231 VMT

A comparative analysis of ECs in the MDA-MB-231 VMT and corresponding VMO scRNA-seq datasets identified pathways that potentially contribute to the observed vascular disruption in the VMT (supplemental figure A4). The ECs from both the VMO and VMT datasets were subset and integrated using scMC [21]. Through unsupervised clustering, three distinct clusters were identified (supplemental figure A4A-A4B): one primarily composed of cells from VMO (EC Normal), another pre-dominantly composed of cells from VMT (EC Tumor), and a third cluster consisting of cells from both datasets (EC Cycling). Pathway analysis conducted on the EC Normal group revealed an increased activity of receptor-ligand interactions and solute carrier family genes associated with the transport of substances across the plasma membrane (supplemental figure A4C). Conversely, the EC Tumor group exhibited elevated matrix metalloproteinase (MMP) activity, as well as cytokine and chemokine activity, suggesting that these ECs may disrupt the surrounding vascular niche (supplemental figure A4D). Lastly, a shared group of cells appeared to be proliferating or cycling (supplemental figure A4E).

ECs from MDA-MB-231 VMT and corresponding VMO were mapped onto a pseudotime using Monocle v2 (supplemental figure A4F). The three EC clusters were further subdivided across five branch points (supplemental figure A4F-A4G). Of particular interest was one branch in the pseudotime trajectory, primarily composed of cells from the tumor (supplemental figure A4F, branch marked with asterisks). To gain deeper insights into the genes influencing this branch and causing its divergence, thereby driving the trajectory split, we performed branch expression analysis modeling (BEAM) using Monocle. For this analysis, branch point two was chosen as the reference for comparing gene expression. BEAM employs two negative binomial general linear models to assess the likelihood of a gene being branch-specific [31]. This approach helps identify genes contributing to a specific branch. The genes associated with the EC tumor branch (see supplemental figure A4H, right-hand side of heatmap) exhibited significant overlap with those identified during pathway analysis, including several MMPs and ADAMTS genes. This finding further highlights the disruption of the vascular niche due to interactions between ECs and MDA-MB-231 TNBC cells.

### Tumor-stromal drug targets were identified by analyzing integrated datasets from VMO to VMT

To further elucidate the transcriptomic shifts between matched VMO and VMT datasets, each pair was integrated with scMC [21] (supplemental figure A5). Integration of MDA-MB-231 VMT and its matched VMO datasets revealed five cell types: ECs, fibroblasts, pericytes, tumor, and cycling cells, as confirmed by the expression of known marker genes (supplemental figure A5A-A5B). In order to identify dysregulated cell-cell communication that might be impacting the vascular niche, we employed LIANA [32] on the individual datasets. LIANA runs multiple cell-cell communication pipelines and compiles the results. The cell-cell communication between MDA-MB-231 VMO and VMT was compared. Results were filtered based on the probability of strong cell-cell communication, revealing that fibroblasts in the VMO communicate with EC through ANGPTL1-TEK (Tie2) signaling. Importantly, this signaling was significantly downregulated in the VMT (supplemental figure A5C). Specifically, fibroblasts from the VMO were found to communicate with EC in the VMO through ANGPTL1-TEK signaling. To validate these interactions, we evaluated the expression of ANGPTL1 and TEK in the integrated dataset, both of which showed decreased expression in the VMT (supplemental figure A5D). The potentially reduced TEK signaling observed in MDA-MB-231 VMT suggests an impairment in vascular maturation and stabilization, as TEK signaling is essential for these processes [33, 34]. And indeed, this is what we saw in the MDA-MB-231 VMTs (Figure 1C).

A similar approach was undertaken for matched HCC1599 datasets, integrating the VMO and VMT datasets, which revealed four cell types: ECs, fibroblasts, pericytes, and tumor cells, as evidenced by the expression of cell type-specific marker genes (supplemental figure A5E-A5F). The cell-cell communication between HCC1599 VMT and VMO was compared, with the filtered results including only interactions unique to the HCC1599 VMT dataset. Notably, one of the top pathways identified was ERBB3/HER3 signaling. To assess the strength of HER3/ERBB3 signaling in both VMO and VMT HCC1599, clusterProfiler [35] was employed (supplemental figure A5G). The analysis revealed that ERBB3 signaling was exclusively present in the VMT dataset, with fibroblasts and tumors acting as ligand sources for the receptor on the tumor population. To test the validity of these findings, we examined the expression of the ligand-receptor pair AREG-ERBB3 (supplemental figure A5H). AREG expression was predominantly found in fibroblasts, with slightly higher expression from cells originating from the VMT. Conversely, ERBB3 expression was limited to cells derived from the VMT, primarily observed in the tumor population. Collectively, these findings support the hypothesis that AREG-ERBB3 signaling may contribute to tumor growth in HCC1599 VMT.

### VMTs recapitulate the tumor microenvironment of clinical TNBC

To validate the clinical utility of the VMT as a tool for identifying therapeutically targetable pathways in TNBC, scRNA-seq datasets derived from primary TNBC samples were integrated with the VMT datasets. Single cell transcriptomic comparison of VMTs with primary samples was conducted to verify whether the ANGPTL1-TEK and AREG-ERBB3 ligand-receptor pair correlations in MDA-MB-231 VMT and HCC1599 VMT and VMOs exhibited comparable patterns of decreased and increased expression, respectively, in clinical specimens. Patient samples from 3 publications[36–38] that included 17 TNBC patients were integrated with both the HCC1599 and MDA-MB-231 VMTs using scMerge [39] (Figure 3). Integration of the datasets resulted in the identification of 12 distinct cell types (Figure 3A), which were consistently observed across most samples (Figure 3B). Cell type identities were assigned based on the expression patterns of established marker genes.

**Fig. 3:**
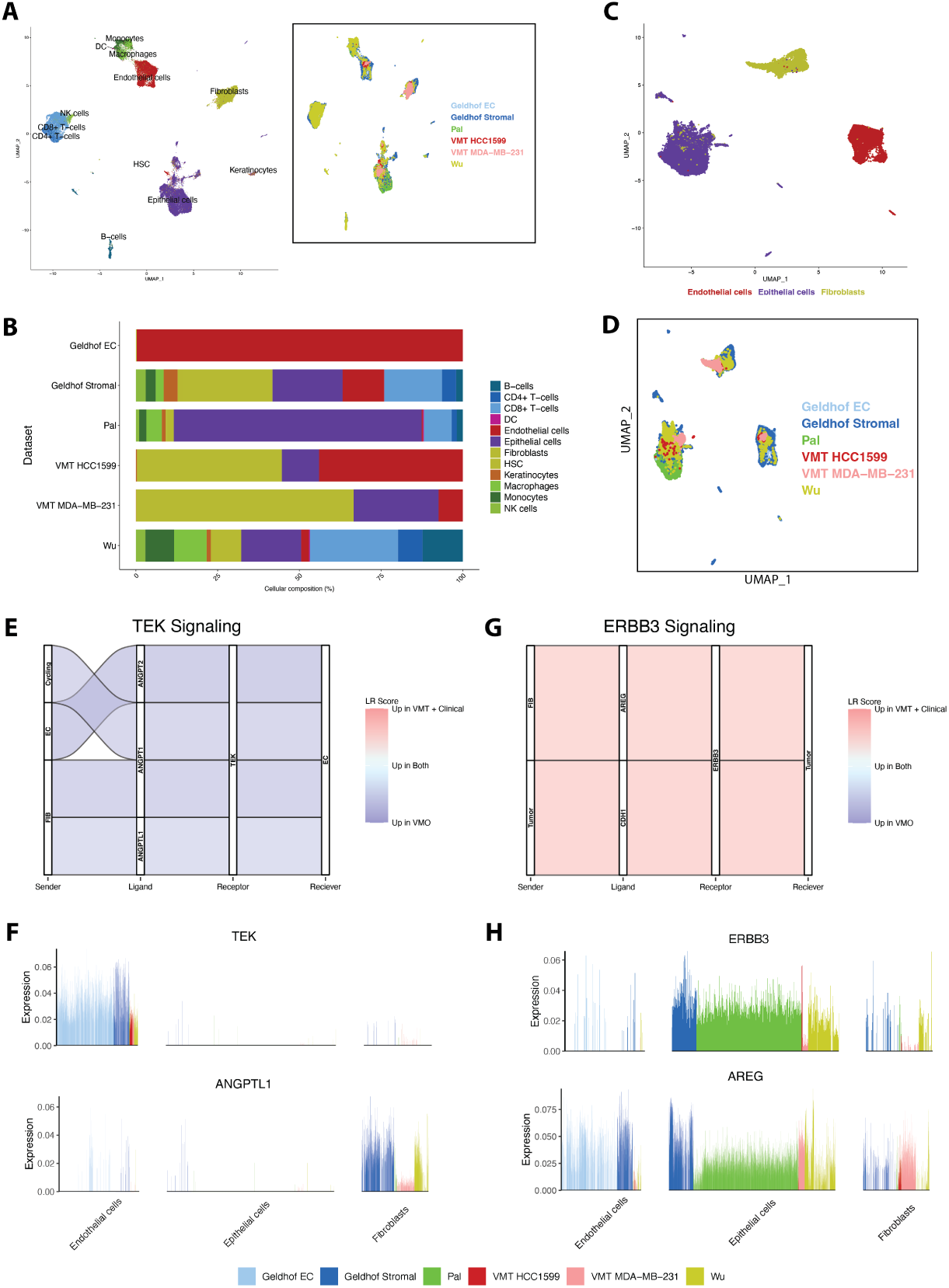
VMTs recapitulate the tumor microenvironment and tumor-stomal signaling pathways of clinical TNBC. **(A)** Integration of VMT HCC1599 and VMT MDA-MB-231, with clinical datasets from three different papers reveals numerous cell types found in patients, including immune cells. Subset shows distribution of datasets in the integrated dimensional space. **(B)** Proportion of each cell type found in each dataset. **(C)** Integration of only the cell types found in the VMT **(D)** shows that the VMT maps well to patient data. **(E)** Angiopoietin-1 receptor (TEK/Tie2) cell-cell communication comparing the strength of the signaling between the VMO and VMT + clinical samples. Sender = cell type that produces the ligand, Receiver = cell type that produces the receptor. **(F)** Erb-B2 Receptor Tyrosine Kinase 3 (ERBB3/HER3) cell-cell communication comparing the strength of the signaling between the VMO and VMT + clinical samples. **(G)** Expression of TEK and TEK ligand angiopoietin like 1 (ANGPTL1) in the integrated dataset. Each bar refers to the expression in a single cell. **(H)** Expression of ERBB3 and ERBB3 ligand amphiregulin (AREG) in the integrated dataset.

To conduct a focused comparison between the TNBC microenvironment of VMTs vs. clinical TNBC, we selectively extracted the fibroblasts, ECs, and epithelial cancer cells from each dataset (Figure 3C). The subset data were then integrated to facilitate a direct comparison between the VMT and primary TNBC datasets. The cells predominantly formed clusters based on their cell type, and cells from each dataset seamlessly integrated with one another (Figure 3D). Cell-cell communication was then performed using LIANA. When comparing the signaling between the clinical specimens and VMTs with the healthy control VMO (MDA-MB-231) dataset, it was observed that ANGPTL1-TEK signaling was more highly expressed in the VMO dataset (Figure 3E), in comparison to either the VMT or the clinical samples, indicating that the expression patterns in the VMTs resembled those found in the clinical specimens. Similarly, when comparing the signaling to the VMO (HCC1599) healthy control, we found that ERRB3 signaling was solely detected in the clinical and VMT integrated dataset (Figure 3G), and not in the corresponding VMO control. The visualization of individual ligand-receptor pairs across datasets allowed us to assess the contribution of each dataset in terms of the cell types producing those transcripts (Figure 3F-3H). For ANGPTL1-TEK, there was a nominal expression of ANGPTL1 in the fibroblast populations that appeared slightly increased compared to the VMTs. For TEK, there seemed to be a somewhat increased expression in the Geldoff EC dataset; however, there was no significant interaction between angiopoietins and TEK. For the AREG and ERBB3 signaling pathway, there was expression of AREG across all the cell types and datasets, supporting its role in signaling to ERBB3, which is expressed exclusively in the tumor epithelial cells. Indeed, targeted Her3 therapeutic approaches may have potential in TNBC [40, 41]. The consistency observed between the VMT and clinical datasets indicates that these therapeutic options show promise for treating patients’ disease.

### Targeting dysregulated Angptl1/Tek tumor-EC signaling in MDA-MB-231 VMT normalizes vessels and improves perfusion

Previous studies have demonstrated the effectiveness of normalizing tumor vasculature in murine mammary carcinoma models using a vascular endothelial protein tyrosine phosphatase (VE-PTP) inhibitor, resulting in improved vessel perfusion and therapeutic delivery to tumors, thereby enhancing tumor killing. VE-PTP is a vascular endothelial cell-specific membrane phosphatase involved in dephosphorylation and consequent inactivation of the Tie2 receptor, leading to vessel destabilization [42]. Razuprotafib (AKB-9778) is a first-in-class VE-PTP inhibitor that induces Tie-2 activation in ECs to promote vascular maturation and enhance tumor perfusion [33]. Given the reduced ANGPTL1/TEK signaling in MDA-MB-231 VMT and clinical TNBC, which correlates with disrupted vasculature, our objective was to investigate the potential benefits of increasing TEK signaling in the tumor EC by blocking VE-PTP with razuprotafib. Thus we looked to integrate a vascular normalization strategy with conventional chemotherapy as a treatment approach for aggressive TNBC.

In order to establish the most effective concentration for treating MDA-MB-231 VMT with razuprotafib, a VE-PTP inhibitor known for its high potency in cell-free assay (IC50 = 17 pM [43]), we conducted a dose-response experiment. Two concentrations, 10 nM and 100 nM, were evaluated in MDA-MB-231 VMT over a 48-hour period, and their vascular normalization capabilities were compared using vascular morphometry analysis, measuring average vessel length, lacunarity (the degree of empty space within the network area), and percent vessel area (Figure 4A). Treatment with 10 nM razuprotafib resulted in a significant increase in average vessel length compared to the control group, along with a significant reduction in vascular lacunarity and a simultaneous increase in the percentage of vascular area within the treated VMTs, consistent with increased vessel density observed with razuprotafib-treated *in vivo* tumors [33]. These results highlight the inherent vascular disruption caused by MDA-MB-231 TNBC in the VMT and demonstrate the possibility of restoring intact vascular networks in the VMT by treatment with razuprotafib. Based on these data we selected the lower concentration (10 nM) of razuprotafib for further experimentation.

**Fig. 4:**
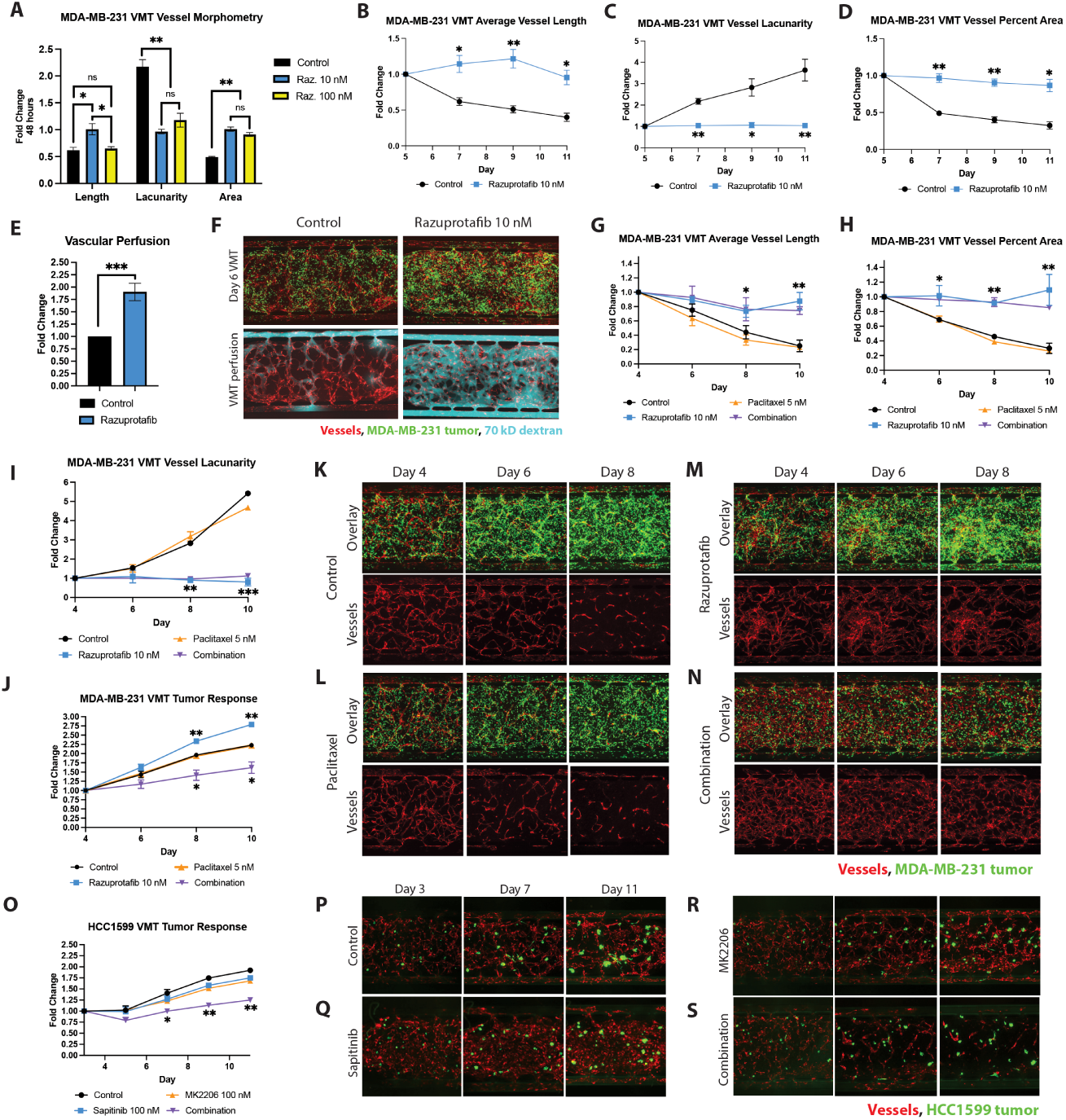
Targeting dysregulated tumor-stromal signaling in MDA-MB-231 VMT and HCC1599 VMT improves therapeutic responses. **(A)** Bar plot showing relative fold change compared to control for different vascular morphometry measurements of MDA-MB-231 TNBC-associated vessel response to razuprotafib 10 nM and 100 nM dose administered in the VMT on day 5 of culture for 48 hours. **(B)** Fold change in average vessel length in MDA-MB-231 VMT over the course of 6 days post-treatment with 10 nM razuprotafib. **(C)** Fold change in vessel lacunarity in MDA-MB-231 VMT over the course of 6 days post-treatment with 10 nM razuprotafib. **(D)** Fold change in vessel percent area in MDA-MB-231 VMT over the course of 6 days post-treatment with 10 nM razuprotafib. **(E)** Fold change in vascular perfusion with 70 kDa fluorescent dextran in MDA-MB-231 VMT 24 hours post-treatment with 10 nM razuprotafib. **(F)** Fluorescent micrographs showing MDA-MB-231 VMT with and without razuprotafib 10 nM, with tumors in green and vessels in red. VMT are perfused on day 6 with 70 kDa fluorescent dextran (shown in cyan). **(G)** Fold change in average vessel length, **(H)** vessel percent area, **(I)** vessel lacunarity, **(J)** and tumor growth in treated MDA-MB-231 VMT. **(K)** Fluorescent micrographs showing control, **(L)** paclitaxel treated, **(M)** razuprotafib treated, **(N)** and combination treated MDA-MB-231 VMT as overlay and vessel only images. **(O)** Fold change in tumor growth in treated HCC1599 VMT. **(P)** Fluorescent micrographs showing control, **(Q)** sapitinib treated, **(R)** MK2206 treated, **(S)** and combination treated HCC1599 VMT.

We proceeded to conduct a time-course experiment in MDA-MB-231 VMT, where a 48-hour treatment of 10 nM razuprotafib was added on day 5. Vascular morphometry analysis was performed every 48 hours for 6 days following the treatment. At each time point examined, the administration of razuprotafib at 10 nM resulted in a significant improvement in average vessel length (Figure 4B), mean vessel lacunarity (Figure 4C), and vessel percent area compared to the control group that persisted for the duration of the experiment (Figure 4D). Within just 24 hours of treatment, there was a substantial (~2-fold) increase in vascular perfusion in MDA-MB-231 VMT compared to the untreated VMT (Figure 4E). These findings are clear when examining fluorescent micrographs of MDA-MB-231 VMTs with and without razuprotafib treatment (Figure 4F), where the pronounced improvement in perfusion of 70 kDa fluorescent dextran in the MDA-MB-231 VMT treated with razuprotafib can be clearly visualized.

### Combining razuprotafib pretreatment with paclitaxel improves chemotherapeutic delivery and anti-cancer response in MDA-MB-231 VMT

In order to assess whether enhanced perfusion would translate into better delivery of therapeutics to the tumor, thereby increasing tumor cell death, pretreatment of MDA-MB-231 VMT with razuprotafib was performed prior to the administration of a dose of paclitaxel that was non-cytotoxic when used as a single-agent therapy in the VMT. Dosing was informed by 2D assay showing marginal effects on MDA-MB-231 tumor growth with 5 nM paclitaxel (supplemental figure A6A). Notably, for each time point examined, there were no significant differences in average vessel length (Figure 4G), vessel percent area (Figure 4H), or vessel lacunarity (Figure 4I) between MDA-MB-231 VMT treated with 5 nM paclitaxel and the control VMT. However, as both single-agent treatment and combination treatment with 5 nM paclitaxel, razuprotafib pretreatment exhibited a statistically significant improvement of the tumor-associated vasculature in all three metrics.

Tumor evaluation revealed that treatment with 5 nM paclitaxel alone did not result in a significant effect on tumor growth compared to the control group. However, when combined with razuprotafib pretreatment, there was a notable improvement in vessel perfusion, leading to enhanced drug delivery and a significant reduction in tumor growth by approximately 30% compared to the control (Figure 4J). Interestingly, the single-agent treatment of razuprotafib resulted in a significant increase in MDA-MB-231 growth, likely due to improved nutrient supply to the tumor. These observations are consistent with the lack of significant effect of razuprotafib on MDA-MB-231 growth observed in 2D cytotoxicity assays either in the absence (supplemental figure A6B) or presence of paclitaxel (supplemental figure A6C).

Examination of the vascular networks in the control MDA-MB-231 VMT revealed progressive deterioration as the tumor grew and invaded the tumor microenvironment (Figure 4K), and a similar disruption of vasculature was observed with paclitaxel treatment as the MDA-MB-231 tumor exhibited approximately a twofold increase in size throughout the duration of the experiment (Figure 4L). In sharp contrast, the administration of razuprotafib resulted in tumor vessel normalization, demonstrated by the preservation of vascular network density despite a significant ~2.75 fold increase in tumor size (Figure 4M). The combination treatment effectively prevented vascular lumen compression and pruning in the tumor microenvironment, facilitating improved drug perfusion and heightened efficacy in tumor eradication (Figure 4N).

### Her3 inhibition synergizes with Akt1/2 inhibition in HCC1599 VMT

Recent clinical trials have highlighted the survival benefits of incorporating AKT inhibition in the treatment of TNBC [44, 45]. Furthermore, studies have shown that the combination of EGFR and Her3 antagonism with AKT inhibition enhances the anti-cancer effects [41, 46]. Considering the activated ERBB3 signaling pathway in HCC1599 VMT, characterized by elevated AREG expression in fibroblasts, and the absence of EGFR expression in HCC1599 [47], we conducted a treatment using sapitinib and MK-2206 to simultaneously inhibit Her3 and Akt1/2, respectively. Sapitinib is an ATP-competitive reversible inhibitor of EGFR/ErbB2/ErbB3, while MK-2206 is an allosteric inhibitor of Akt1/2.

As illustrated in Figure 4O-4S, when HCC1599 VMT was treated with 100 nM sapitinib and 100 nM MK-2206 individually, no significant cytotoxic activity against HCC1599 TNBC cells was observed. However, when combined, these treatments synergized to suppress tumor growth in the VMT, resulting in approximately 75% reduction in tumor burden compared to control and single-agent treatments. In contrast, HCC1599 exhibited no sensitivity to sapitinib, MK-2206, or the combination treatment in 2D culture (see supplementary figure A6D-A6G) [48]. The insensitivity observed in 2D monocultures is likely attributable to the absence of paracrine signaling to ErbB3, which, in contrast, is captured within the VMT.

## 3 Discussion

To study the TNBC TME, we established vascularized microtumors (VMTs) using two well-studied TNBC cell lines, MDA-MB-231 and HCC1599, along with ECs and primary stromal cells derived from histopathologically normal human breast tissue. MDA-MB-231 is an aggressive and poorly differentiated claudin-low molecular subtype TNBC cell line, known for its invasiveness and metastatic potential in various organs of mice [49]. Our findings demonstrate that the growth and migration of MDA-MB-231 cells in the VMT mirrors that seen *in vivo* (highly invasive) and leads to significant disruption of the vasculature, characterized by vessel regression, reduced vascular density, and impaired perfusion. These vascular changes observed in the VMT closely resemble the structural and functional vascular disruption seen in solid tumors, which often contributes to tumor progression by hindering effective delivery of anti-tumor therapies [8, 50, 51].

By analyzing the TNBC VMT compared to matched vascularized micro-organs (VMO), our aim was to uncover the molecular changes occurring in normal breast-tissue derived stromal cells when exposed to the TNBC microenvironment. Through single-cell transcriptomic analyses, we identified distinct cell-cell communication patterns and gene expression signatures within the tumor-stromal niche of MDA-MB-231. Notably, populations of fibroblasts exhibited gene signatures associated with cancer-associated fibroblasts (CAFs) as well as synthetic fibroblasts. The enriched fibroblast populations exhibited altered extracellular matrix organization processes that likely contributed to the destabilization of the tumor-associated vasculature. These characteristics observed within the tumor microenvironment have been previously associated with unfavorable outcomes in TNBC [11], underscoring the potential of utilizing the VMT to gain deeper insights into the intricate interplay within the TNBC TME.

Putative targets from cell-cell communication analyses were further validated for clinical relevance by comparing scRNA-seq data from VMTs/VMOs with publicly available scRNA-seq datasets derived from primary TNBC specimens. Significantly, the dysregulated ANGPTL1/TEK signaling pathway emerged as an intriguing therapeutic target with potential clinical significance since signaling through this ligand-receptor pair was found to be present in the VMO (MDA-MB-231) but reduced in the MDA-MB-231 VMT and TNBC clinical specimens. TEK, also known as Tie-2, is an endothelial tyrosine kinase receptor involved in vascular stabilization that is activated by angiopoietins, including Angiopoeitin-related protein 1 (ANGPTL1) [42]. Its activity is regulated by VE-PTP, which dephosphorylates and inactivates it. Notably, in mouse models the VE-PTP inhibitor AKB-9778, or razuprotafib, has shown promise in stabilizing breast cancer vasculature and suppressing metastatic progression by sustaining Tie-2 activation [33]. Our findings using a human cell-based platform demonstrate that pharmacological inhibition of VE-PTP in the VMT can restore the structure and function of tumor vessels, leading to increased sensitivity of TNBC to low-dose chemotherapy with paclitaxel. However, it should be noted that as of the publication date, no clinical trials have been conducted to evaluate the effectiveness of razuprotafib in combination with anti-cancer agents, specifically in TNBC or any other cancer type [52]. This presents a promising opportunity for future clinical trials in TNBC, exploring the use of razuprotafib as a pretreatment strategy to enhance the delivery of cytotoxic drugs and immune cells to the tumor, ultimately leading to improved patient outcomes.

In our study, we observed that TNBC cells in both HCC1599 VMT and clinical specimens upregulate Her3 (ERBB3) in response to paracrine signaling from fibroblasts, mainly through increased expression of the ligand amphiregulin (AREG). HCC1599, a basal-like TNBC cell line, has been shown to better represent primary TNBC disease based on various “-omics” metrics compared to other cell lines [18, 19]. It also carries a BRCA2 gene mutation [47]. Interestingly, despite Her3 and Akt inhibitors showing little efficacy in HCC1599 monocultures as single agents or in combination, testing a low-dose combination of sapitinib and MK2206 in the HCC1599 VMT revealed synergistic anti-cancer effects. Sapitinib is a reversible ATP-competitive inhibitor of ErbB1/2/3 with an IC50 of 4 nM for ErbB3 in cell-free assays. Since HCC1599 lacks Her2 and EGFR expression [47], sapitinib presumably acts through ErbB3 inhibition in the VMT.

Previous phase II trials of sapitinib in combination with paclitaxel for advanced breast cancer did not meet their primary endpoint of significantly increased duration of progression-free survival in the combination arm [53]. Similarly, a phase II trial evaluating MK2206-2HCl, a highly selective inhibitor of Akt1/2 [54], as a monotherapy showed minimal therapeutic activity. However, a similar clinical trial assessing the combination of Akt inhibition as combination therapy with paclitaxel did show promising results with increased progression-free survival [45]. Notably, recent preclinical studies have highlighted the importance of targeting both Her3/EGFR and the downstream mediator Akt for optimal response in TNBC, particularly in TNBC cells with mutations in PIK3CA, AKT, or loss of PTEN [46, 55]. Although HCC1599 lacks these mutations, it does harbor mutations in PI3K/Akt pathway mediators, such as TNS1 and PIK3C2B [47], which may also play a role in predicting patient’s response.

By faithfully reproducing essential elements of the tumor microenvironment, such as niche factors, dynamic cell-cell interactions, and the intricate tumor structure consisting of cancer cells, stromal cells, and vasculature, the VMT provides remarkable possibilities for modeling cancer within a physiologically relevant context. Recently, federal organizations, academic institutions, and industry experts have also recognized the value of tumor chips in expediting oncology drug discovery and streamlining preclinical development processes [56]. Our study further underscores the importance of faithfully reproducing the specific characteristics of the TNBC microenvironment to facilitate the discovery of innovative therapeutic strategies targeting reprogrammed stromal components. These efforts will be invaluable in accelerating the translation of findings from preclinical research into clinical trials, ultimately benefiting patients diagnosed with TNBC.

## 4 Conclusion

Through analysis of an advanced VMT model, clinically relevant drug targets associated with tumor-stromal interactions in TNBC were identified. The VMT faithfully reproduces critical components of the tumor microenvironment, encompassing tumor, stromal cells, and vasculature within a complex tissue structure. As such, the VMT can serve as a powerful tool for the research community and may also be useful for personalized medicine applications. In this study, effective reduction of TNBC growth in the HCC1599 VMT model was achieved by targeting the activated AREG/ERBB3 signaling pathway, which is also activated in clinical TNBC. Similarly, in the MDA-MB-231 VMT model, targeting the dysregulated ANGPTL1/TEK signaling, also observed in clinical TNBC, resulted in the normalization of dysfunctional vasculature and improved paclitaxel delivery, leading to reduced TNBC growth. These findings highlight the potential of modulating tumor-stromal crosstalk as a promising therapeutic strategy for TNBC. Finally, we note that the VMT platform is also ideally suited to studies of immune cell-based approaches to tumor management.

## 5 Methods

### Microfluidic device fabrication

Device fabrication and loading has been described previously [13, 14, 16, 17, 57]. Briefly, a customized polyurethane master mold is fabricated using 2-part polyurethane liquid plastic (Smooth Cast 310, Smooth-On Inc.). A PDMS layer is then replicated from the master mold, holes are punched for inlets and outlets, and the platform is assembled in two steps. The PDMS layer is first attached to the bottom of a 96-well plate by chemical gluing and oxygen plasma treatment for 2 minutes. A 150 µm thin transparent membrane is then bonded to the bottom of the PDMS device layer by treating with oxygen plasma for an additional 2 minutes. The fully assembled platform is placed in 60 *^◦^*C oven overnight, covered with a standard 96-well plate polystyrene lid, and sterilized using UV light for 30 minutes prior to loading cells.

### Collection and processing of primary human normal breast tissue

A breast tissue sample was acquired after ethical approval by the research center’s Institutional Review Board (IRB) from the Cooperative Human Tissue Network (CHTN). The patient gave written, informed consent and shared the respective metadata. Inclusion criteria was that the sample was histopathologically normal. Tissue was processed as previously reported in [58]. The surgical specimen was washed in PBS, mechanically dissociated with scalpels, digested with 2 mg*/*mL collagenase I (Life Technologies, 17100-017) in DMEM (Corning, 10-013-CV) overnight, digested in 20 U*/*mL DNase I (Sigma-Aldrich, D4263-5VL) for 5 minutes, and centrifuged for 2 minutes at 150*×*g. From the healthy patient specimen, supernatant was collected and centrifuged for 5 min at 500*×*g to isolate epithelial tissue chunks in the pellet. These were viably cryopreserved in DMEM with 50% FBS (Omega Scientific, FB-12) and 10% DMSO (vol/vol) before processing into single cells for flow cytometry. For flow cytometry cells were stained for FACS using fluorescently labeled antibodies for CD31 (eBiosciences, 48-0319-42), CD45 (eBiosciences, 48-9459-42), EpCAM (eBiosciences, 50-9326-42), CD49f (eBiosciences, 12-0495-82), SytoxBlue (Life Technologies, S34857), PROCR (Biolegend, 351904), and PDPN (BioLegend, 337014).

### Cell culture and microfluidic device loading

To load cells, primary human normal breast stromal cells and ECFC-ECs were harvested and resuspended in fibrinogen solution at a concentration of 7 *×* 10^6^ cells*/*mL and 3*×*10^6^ cells*/*mL, respectively. MDA-MB-231 and HCC1599 TNBC cells were introduced at a concentration of 1 *×* 10^5^ to 2 *×* 10^5^ cells*/*mL to establish VMT. Fibrinogen solution was prepared by dissolving 70% clottable bovine fibrinogen (Sigma-Aldrich) in EBM2 basal media (Lonza) to a final concentration of 5 mg*/*mL. The cell-matrix suspension was mixed with thrombin (50 U*/*mL, Sigma-Aldrich) at a concentration of 3 U*/*mL, quickly seeded into the microtissue chambers, and allowed to polymerize in a 37 *^◦^*C incubator for 15 minutes. Laminin (1 mg*/*mL, LifeTechnologies) was then introduced into the microfluidic channels through medium inlets and incubated at 37 *^◦^*C for an additional 15 minutes. After incubation, culture medium (EGM-2, Lonza) was placed into the microfluidic channels and medium wells. Medium was changed every other day and hydrostatic pressure head re-established every day to maintain interstitial flow.

### XTT assay for 2D cytotoxicity

Cell viability in 2D monolayer cultures ± treatment was quantified using an XTT assay according to the manufacturer’s protocol (Sigma-Aldrich). Briefly, 5,000 MDA-MB-231 or HCC1599 TNBC cells were seeded in triplicate in a 96-well plate and allowed to grow for 8 hours prior to treatment with increasing doses of paclitaxel, razuprotafib, sapitinib, MK-2206 or combination treatments. XTT assays were performed after 48 hours of drug exposure. A standard cell dilution was used to quantify total cell numbers. Cell viability was normalized to control wells without drug treatment.

### Drug treatment in the VMT

Hydrostatic pressure was restored daily in the high throughput platform, while medium was changed every two days. After culturing for 4–5 days to allow a perfused vasculature to form within each VMT, culture medium was replaced with medium containing the drugs at the desired concentration. Drugs were delivered to the tumor through the vascular bed via gravity-driven flow. Sapitinib (reversible, ATP-competitive inhibitor of EGFR, ErbB2 and ErbB3), MK-2206 2HCl (Akt 1/2/3 inhibitor), and paclitaxel (microtubule stabilizer) were purchased from SelleckChem.

Razuprotafib is a VE-PTP inhibitor that was purchased from MedChemExpress. MDA-MB-231 VMT were randomly assigned to one of four conditions: control (vehicle only), 5 nM paclitaxel, 10 nM razuprotafib, or combination 5 nM paclitaxel with 10 nM razuprotafib. VMT receiving razuprotafib were treated on day 4 for 24 hours, followed by 48 hour paclitaxel treatment for the combination condition or complete media for the single-agent condition. Single-agent paclitaxel was also treated for 48 hours. HCC1599 VMT were treated on day 4-5 with one of the following: control (vehicle only), 100 nM sapitinib, 100 nM MK-2206 2HCl, or combination 100 nM sapitinib with 100 nM MK-2206 2HCl. Complete medium was replaced after 48 hours. Fluorescent micrographs of VMT were taken every 48 hours for 6 days post-treatment and growth of the tumor was quantified.

### Fluorescence imaging and analyses

Fluorescence images were acquired with a Biotek Lionheart fluorescent inverted microscope using automated acquisition and standard 10x air objective. AngioTool software (National Cancer Institute) was used to quantify vessel area, vessel length, number of vascular junctions and endpoints in the VMT. ImageJ software (National Institutes of Health) was utilized to measure vessel diameter and measure the total fluorescence intensity (i.e. mean grey value) for each tumor image to quantify tumor growth. Each chamber was normalized to baseline. Vessels were perfused by adding 25 µg/mL FITC-or rhodamine-conjugated 70 kDa dextran to the medium inlet. Once the fluorescent dextran had reached the vascular network, time-lapse image sequences were acquired using a Nikon Ti-E Eclipse epifluorescence microscope with a 4× Plan Apochromat Lambda objective. Perfusion images were analyzed using ImageJ software by measuring change in fluorescence intensity within perfusable vessel regions and creating a composite score based on total perfusable vascular area. Tumor growth in the VMTs was quantified by measuring the total fluorescence intensity in the color channel representing the tumor cells. This takes into account both the area of the individual tumors and the depth, as thicker areas appear brighter. Any adjustments made to images are performed on the entire image, with all images in that experimental group adjusted to the same settings.

### Cell harvesting and single-cell sequencing

Media was aspirated from each well and the high-throughput plates were inverted so that the device layer was facing up. The bottom membrane was gently peeled off the device to expose the tissue chambers. Devices were first washed with 500 µL of HBSS. To each device unit, 100 µL of TrypLE was added and allowed to sit as a bubble on top of the device while the plate was put back in the 37 *^◦^*C incubator for 5 minutes. After digestion, tissue chambers were pipet into a 15 mL conical with EGM2 to neutralize the TrypLE. Digestion solution containing the dislodged tissues was centrifuged at 300*×*g for 5 minutes at 4 *^◦^*C to pellet single cells and whole tissues. Media was carefully aspirated and a solution of 1 mg*/*mL (200 U/mL) collagenase III in HBSS was added to the undigested tissues. After gentle pipetting, the solution was allowed to sit for 2 minutes at room temperature before pipetting gently again to dissociate the gel. Digestion mix was washed with 10 mL of EGM2 and centrifuged at 300*×*g for 5 minutes at 4 *^◦^*C. Cells were resuspended into 1X DPBS with 1% human serum albumin and passed through a pre-wetted 70 µm filter by spinning at 200*×*g for 1 minute. Cells were counted and volume was adjusted so that total cells were at a concentration of 1000 cells*/*µL. Cellular suspensions were loaded onto a Chromium Single Cell Instrument (10X Genomics) to generate single-cell gel beads in emulsion (GEMs). GEMs were processed to generate cDNA libraries by using 10X Genomics v2 chemistry according to the Chromium Single Cell 3’ Reagents kits v2 user guide: CG00052 Rev B. Quantification of cDNA libraries was performed using Qubit dsDNA HS Assay kit (Life Technologies Q32851), high-sensitivity DNA chips (Agilent 5067-4626) and KAPA qPCR (Kapa Biosystems KK4824). Libraries were sequenced on an Illumina NovaSeq6000 to achieve an average of 50,000 reads per cell.

### Transcriptome alignment and data processing of VMO and VMT libraries

FASTQ files for each library were aligned to an indexed GRCh38 reference genome using Cell Ranger (10X genomics) Count 3.1.0 to generate count matrices. Count matrices were loaded into RStudio (1.2.5042 version, R version 4.0.4 [59, 60]) and converted into a Seurat object using the Seurat R Package (version 4.0.0.9015) [61, 62]. Cells containing more than 15% mitochondrial RNA or with less than 200 features were removed. Additionally, genes not detected in at least 3 cells were trimmed. Data were normalized, the 2000 top variable features were found, and scaled using the Nor-malizeData, FindVariableFeatures, and ScaleData functions, respectively. Data were further normalized using the SCTransform function and the difference between cells in the S phase and G2M phase was regressed to help distinguish non-cycling from cycling cells. The ‘SCT’ assay slot was used for analyzing the individual datasets. Doublets were identified and removed using the DoubletFinder R package (version 2.0.3) [63]. A modified version of the chooseR R package [64] was used to generate silhouette scores for selecting the resolution. Clusters were labeled based on the expression of known markers [ECs: PECAM1, CDH5, VWF, KDR, CLDN5 [65], Fibroblasts: PDGFRA, PDGFRB, COL1A1 [66], Stroma: PDGFRB, TPM1, ACTA2 [66], Cycling cells: MKI67 and TOP2A [67], and Tumor: KRT18 and EPCAM [68]], and cell types were later confirmed using the SingleR R package (version 1.4.1) [69]. Heatmaps were generated by filtering expression for each clustering by log2 fold change. Pathway analysis was performed using clusterProfiler [35] with the GO Biological Process 2021 database on the top 100 differentially expressed genes. The significance of key pathways was directly compared for each dataset.

### Integration between VMO and VMT libraries

Individual libraries were integrated using the scMC RunscMC R function and package (version 1.0.0) with default settings [21]. scMC removes the technical variation while preserving biological variation using variance analysis in an unsupervised manner. For integration the ‘RNA’ slot was used for each library. After integration, cell types were assigned to individual cells based on the analysis of the individual libraries. UMAP for all of the datasets was generated with palette from the Polychrome R package (version 1.5.1)[70].

### Processing publicly available patient libraries and integrating with VMT libraries

The Geldhof [38] libraries contain 8,433 ECs from 9 breast cancer patients and 26,515 total cells (GSE155109). The EC enriched and stromal population count matrices were downloaded from the supplementary files. The Pal [37] libraries contain four libraries from triple negative tumor with additional microenvironment cells (GSM4909284, GSM4909283, GSM4909282, and GSM4909281). The Wu [36] libraries contain 5 patient triple-negative breast cancer libraries that total 600 ECs (Broad SCP1106). The normalization and filtering of cells was performed as previously described. The clustree R package [71] was used to determine the optimal resolutions. The cells were assigned labels using the SingleR package, with the reference database limited to only include cell types that were mentioned in the original papers. The publicly available libraries and the VMT libraries were integrated with the scMerge (version 1.9.99) [39] R package using the scMerge::scMerge2 [72] with stably expressed genes calculated for the dataset to normalize the count matrices. Labels were assigned to each cell type using SingleR and 82 cells were removed due to not being assigned a label and not having a consensus cell type as its nearest neighbor. To integrate only the cell types present in the VMT libraries, the endothelial cells, epithelial cells, and fibroblasts as labeled by SingleR for each dataset were subset, and the subset libraries were integrated as described for the full libraries.

### Cell-cell communication, visualization, and bar plot generation

The liana (version 0.0.1) [32] R package was used to compute the cell-cell communication between cell types. Possible interactions were pulled from the OmniPath database [73]. Liana aggregates results from multiple cell-cell communication pipelines, and the results from the CellChat [74], Conectome [75], iTALK [76], NATMI [77], and SingleCellSignalR [78] pipelines were computed. Interactions that had a p-value *<*0.05 for CellChat were subset. Interactions were further filtered based on an aggregate rank *<*0.05. Significant interactions were filtered based on expression in the VMO or VMT datasets. Individual receptor interactions were visualized using the CrossTalkeR R package (version 1.3.2)[79].

### Pseudotime

Pseudotime was performed using Monocle v2 [31, 80]. Seurat data from the RNA slot was converted into a CellDataSet format following custom code provided by UC Irvine’s Genomic High Throughput Facility. The expression for the data was modeled using a negative binomial distribution with the monocle::newCellDataSet function with the lower detection limit set to 0.5. The size factors and dispersion values that help with normalization and differential expression were computed, low-quality cells were filtered out, and cells were assigned to a pseudotime trajectory based on the total amount of transcriptional change. Branch expression analysis modeling (BEAM) was run to identify the branch-dependent genes. BEAM generates two negative binomial general linear models, one assuming the gene is branch independent and a second assuming the gene is branch dependent. The models are then fit to the pseudotime branch trajectories, and a likelihood ratio test determines if the genes are branch dependent or independent. Finally, a heatmap with the branch-dependent genes was generated using the monocle::plot genes branched heatmap function. Genes with less than a 1e-45 q-value (78 genes) were shown on the heatmap.

### Statistical analyses

Data are represented as mean ± standard error. Comparison between experimental groups of equal variance were analyzed using an unpaired t-test and 95% confidence interval or one-way ANOVA followed by Dunnett’s test for multiple comparisons. Statistical calculations were performed using GraphPad Prism 9.0 (GraphPad Software, San Diego, California USA, www.graphpad.com).

## Supplementary information

Supplemental table 1. Pathway analysis for VMOs and VMTs (both MDA-MB-231 and HCC1599) by cell type.

Supplemental table 2. Top 100 differentially expressed genes for fibroblast populations.

### Abbreviations

CAF: cancer-associated fibroblast
EC: endothelial cell
ECM: extracellular matrix
TME: tumor microenvironment
TNBC: triple negative breast cancer
VMO: vascularized micro-organ
VMT: vascularized micro-tumor

## Declarations

## Acknowledgments

We would like to thank Daniel Kim for his assistance in the lab. This work was made possible, in part, through access to the following: the Genomics Research and Technology Hub (formerly Genomics High-Throughput Facility) Shared Resource of the Cancer Center Support Grant (P30CA-062203), the Single Cell Analysis Core shared resource of Complexity, Cooperation and Community in Cancer (U54CA217378), the Genomics-Bioinformatics Core of the Skin Biology Resource Based Center @ UCI (P30AR075047) at the University of California, Irvine and NIH shared instrumentation grants 1S10RR025496-01, 1S10OD010794-01, and 1S10OD021718-01.

## Availability of data and materials

Single cell RNA sequencing data generated in this study has been deposited to the GEO database (cite number once deposited).

### Ethics approval and consent to participate

A breast tissue sample was acquired after ethical approval by the research center’s Institutional Review Board (IRB) from the Cooperative Human Tissue Network (CHTN). The patient gave written, informed consent and shared the respective metadata.

### Consent for publication

All authors have reviewed and provided their consent for the publication of the content presented in this manuscript.

## Authors’ contributions

SJH conceived of the study idea, performed the majority of experiments, and wrote the manuscript; CJH provided intellectual input, performed the majority of analyses, and wrote the manuscript; DG provided intellectual input, performed XTT assay and analysis, and edited the manuscript; AM processed and measured fluorescent micrographs and edited the manuscript; KN supplied primary breast tissue-derived stroma, provided intellectual input, and edited the manuscript; KK provided intellectual input and edited the manuscript; CCWH provided intellectual input, manuscript editing, and overall project management.

## Appendix A Supplementary materials

**Fig. A1:**
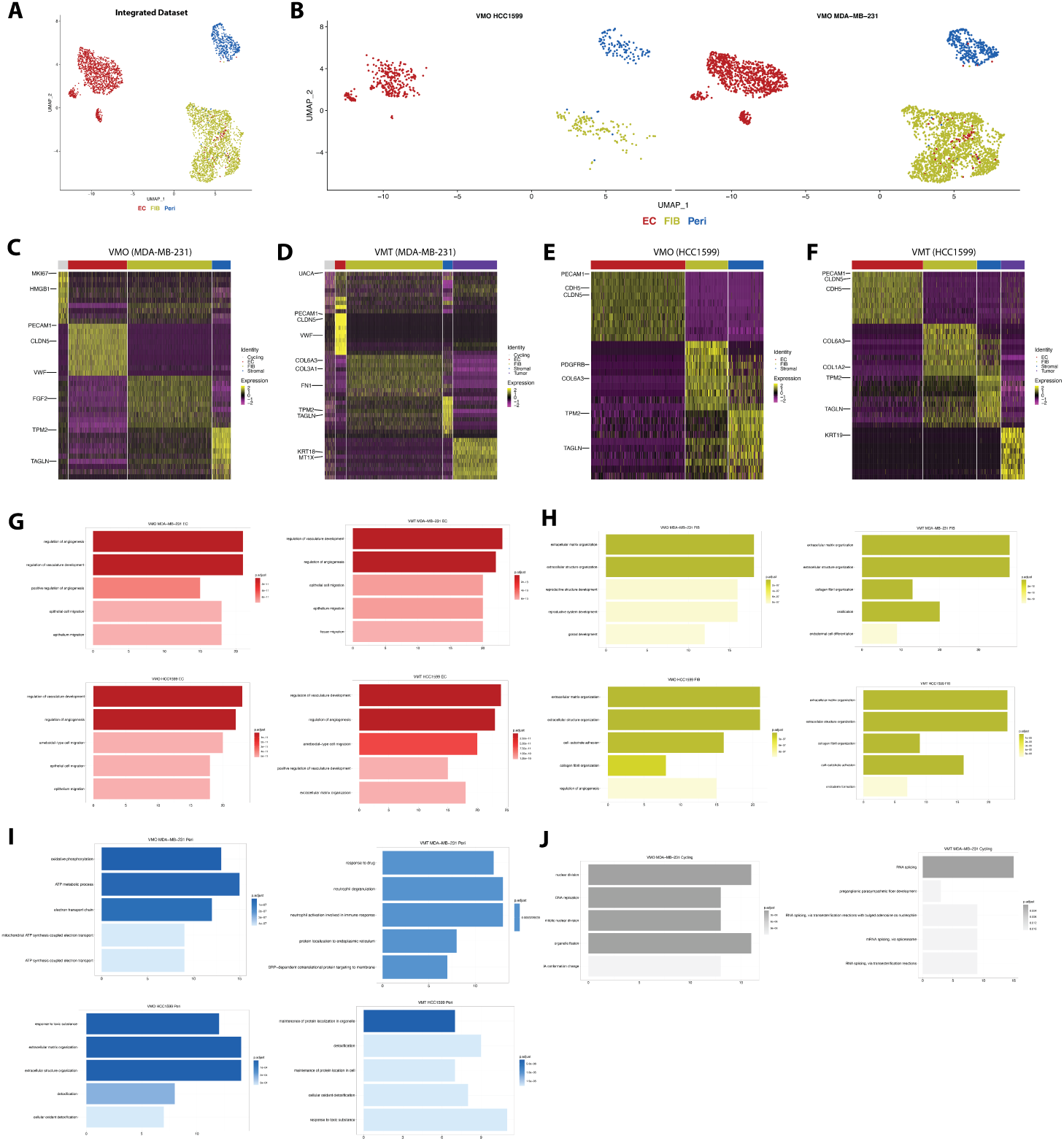
Differential gene expression analyses of individual datasets reveal distinct pathway activation dependent on cell type in both VMTs and VMOs. **(A)** UMAP showing integrated VMO dataset, **(B)** shown with datasets split by VMO matched to HCC1599 and MDA-MB-231 VMT, respectively. **(C)** Heatmap showing top 10 DEGs for VMO matched to MDA-MB-231 VMT. **(D)** Heatmap showing top 10 DEGs for MDA-MB-231 VMT. **(E)** Heatmap showing top 10 DEGs for VMO matched to HCC1599 VMT. **(F)** Heatmap showing top 10 DEGs for HCC1599 VMT. **(G)** Gene Ontology Biological Processes pathway analyses for endothelial cells (EC) in each dataset. (H) Gene Ontology Biological Processes pathway analyses for fibroblasts in each dataset. **(I)** Gene Ontology Biological Processes pathway analyses for pericytes in each dataset. **(J)** Gene Ontology Biological Processes pathway analyses for cycling cell population in each dataset.

**Fig. A2:**
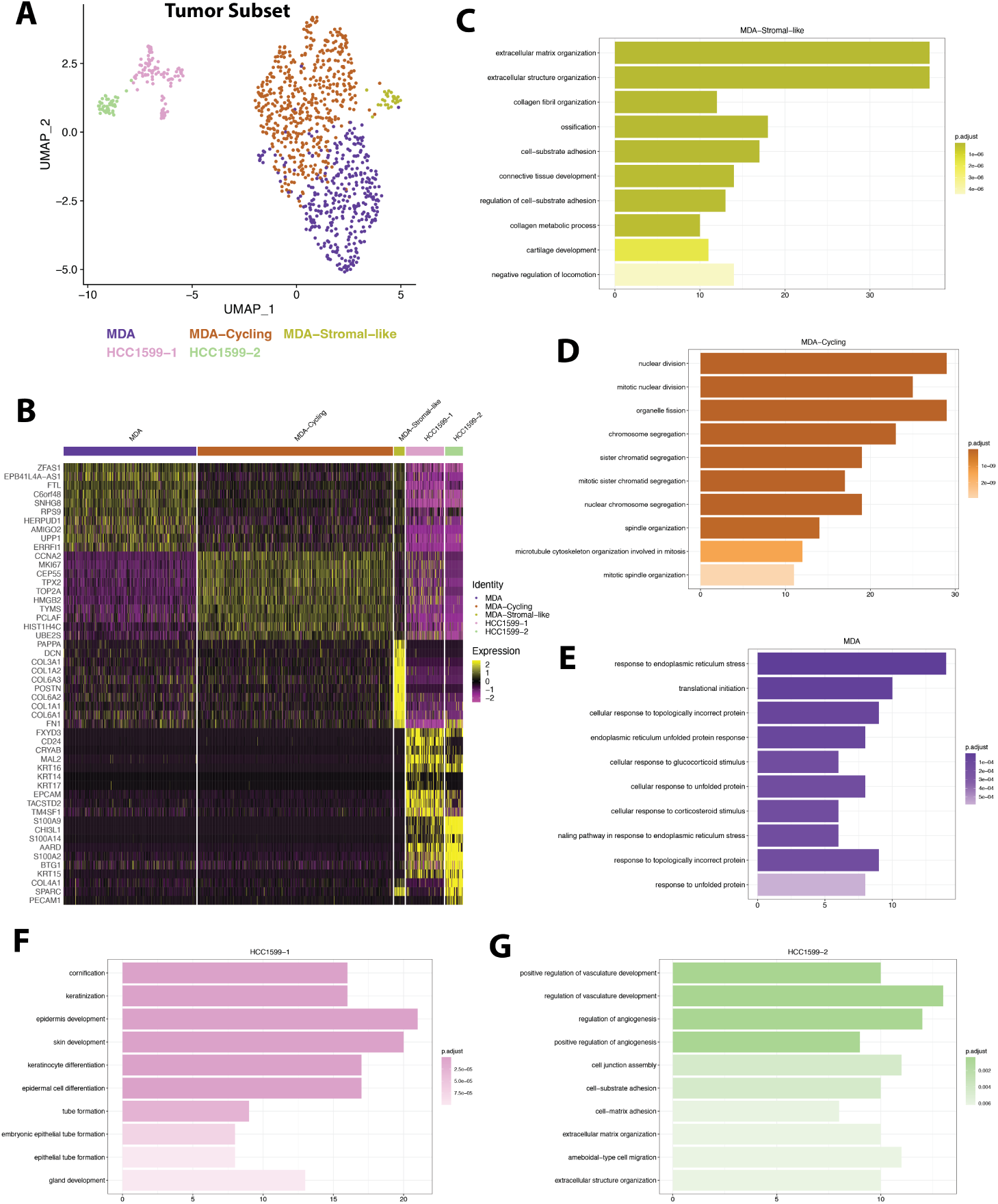
Tumor subset from integrated VMT dataset highlights differences in TNBC cell populations. **(A)** UMAP showing cell clusters for integrated VMT dataset. **(B)** Heatmap showing top 10 DEGs for tumor populations from MDA-MB-231 and HCC1599 VMTs. **(C)** Gene Ontology Biological Processes pathway analyses for MDA-MB-231 stromal-like tumor population, **(D)** MDA-MB-231 cycling tumor population, and **(E)** MDA-MB-231 tumor population. **(F)** Gene Ontology Biological Processes pathway analyses for HCC1599-1 tumor population and **(G)** HCC1599-2 tumor population.

**Fig. A3:**
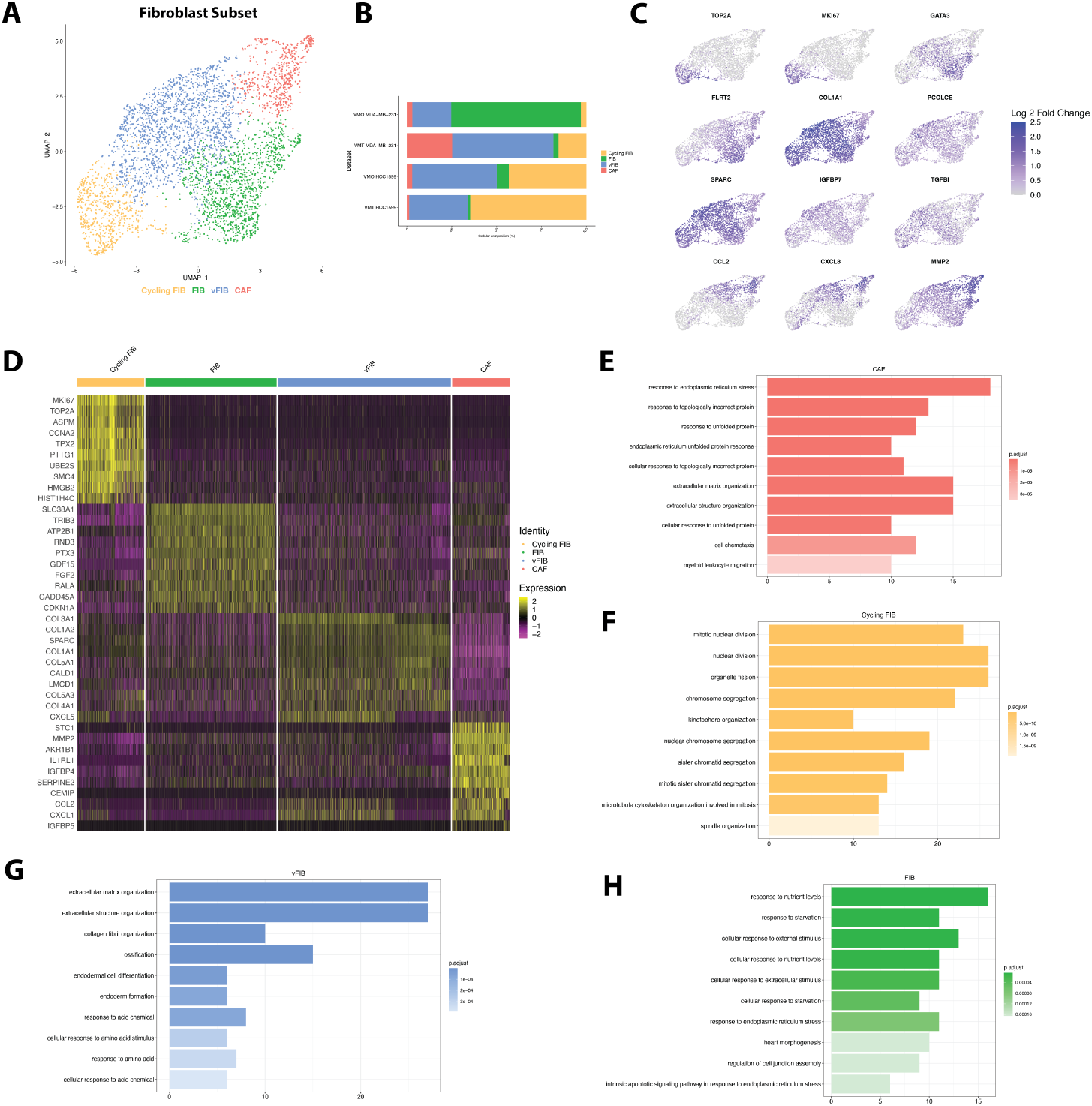
Fibroblast subset from fully integrated dataset shows increase in CAF cell signature in MDA-MB-231 VMT. **(A)** Unsupervised clustering of integrated fibrob-last dataset reveals four distinct clusters with **(B)** various proportions of each dataset in the cluster. **(C)** Expression profiles of differentially expressed genes. **(D)** Heatmap of top 10 differentially expressed genes (DEGs) for each cluster. **(E)** Pathway analysis for the top 100 DEGs for each cluster for the CAFs, **(F)** cycling fibroblasts, **(G)** synthetic fibroblasts, and **(H)** fibroblasts.

**Fig. A4:**
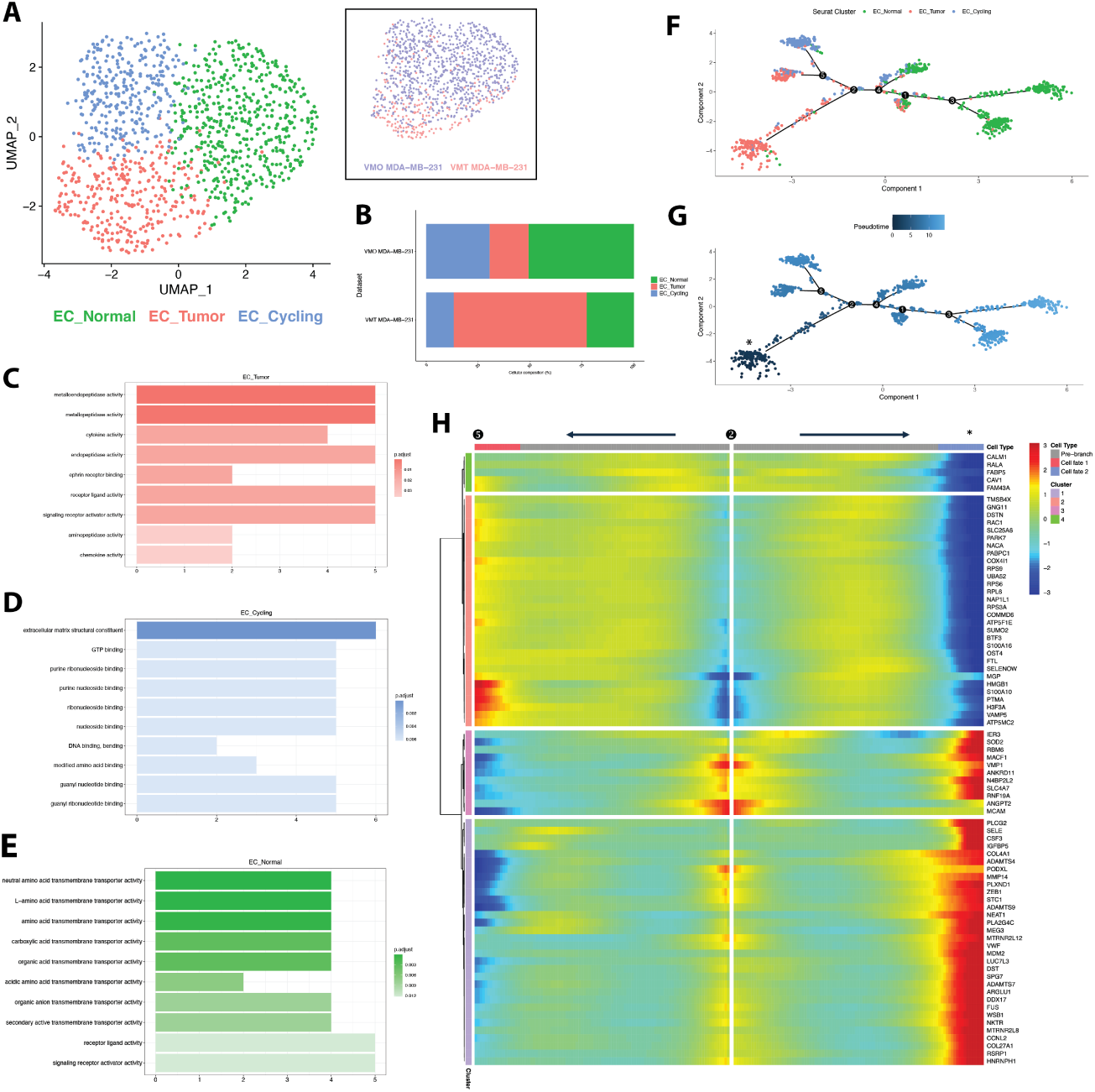
EC subset pseudotime analysis reveals enrichment of tumor-associated EC signature in MDA-MB-231 VMT. **(A)** Integration and unsupervised clustering of the endothelial cells (ECs) from the VMO and VMT MDA-MB-231 reveal three populations. Subset: the distribution of cells based on dataset in the dimensional space. **(B)** The proportion of each cluster from each dataset. Note the distinction between the normal and tumor populations. **(C)** Pathway analysis on the top 30 differentially expressed genes for each cluster shows the normal EC with an increase in receptor-ligand activity and transmembrane transporter activity, **(D)** the tumor EC have an increase in metalloendopeptidase activity and cytokine activity, and **(E)** the cycling EC have an increase in DNA binding. **(F)** Pseudotime on the MDA-MB-231 ECs supports the prior clustering and reveals a branch off branch point 2 that is mainly tumor associated (denoted with *). **(G)** Assigning pseudotime emphasizes differences between branch points and shows a starting time with normal EC transitioning into tumor EC. **(H)** The heatmap displays the expression of genes associated with specific branches. Branch 2 is positioned at the center of the heatmap, serving as the midline. Moving towards the left from the center indicates gene expression linked to cells at branch point 5. Moving towards the right indicates gene expression associated with tumor ECs.

**Fig. A5:**
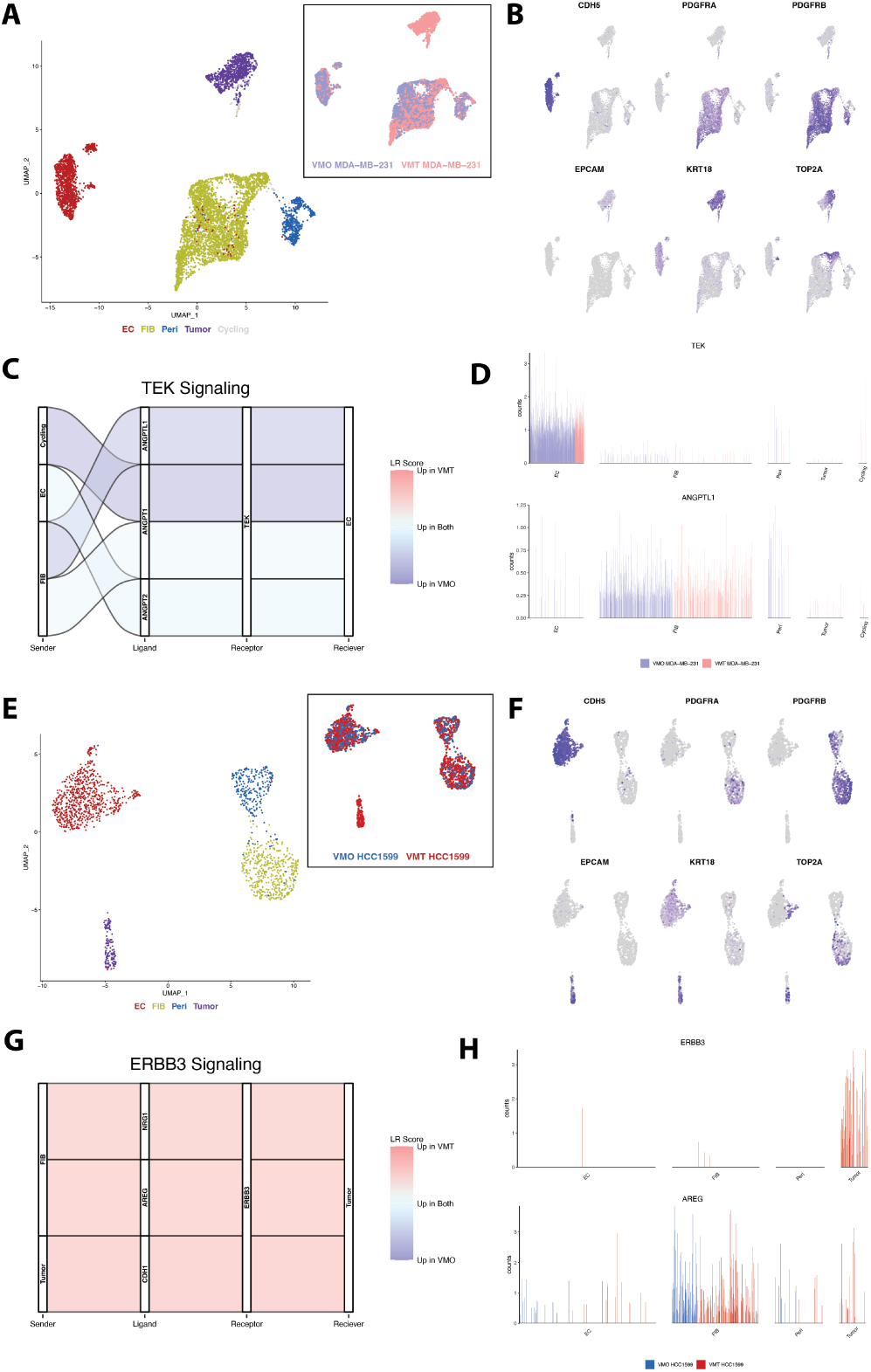
Tumor-stromal drug targets are identified in VMO to VMT integrated datasets. **(A)** Integration with the VMO MDA-MB-231 and VMT MDA-MB-231 datasets showing five cell types of cycling cells, ECs, fibroblasts (FIB), stromal cells (Peri), and tumor **(B)** as demonstrated by the expression of key marker genes. **(C)** Angiopoietin-1 receptor (TEK) cell-cell communication comparing the strength of the signaling between the VMO and VMT. Sender = cell type that produces the ligand, Receiver = cell type that produces the receptor. **(D)** Expression of TEK ligand angiopoietin like 1 (ANGPTL1) and TEK in the integrated dataset. Each bar refers to the expression in a single cell. **(E)** Integration with the VMO HCC1599 and VMT HCC1599 datasets showing four cell types of endothelial cells (ECs), fibroblasts (FIB), pericytes (Peri), and tumor **(F)** as demonstrated by the expression of key marker genes. **(G)** Erb-B2 Receptor Tyrosine Kinase 3 (ERBB3/HER3) cell-cell communication comparing the strength of the signaling between the VMO and VMT. **(H)** Expression of ERBB3 ligand amphiregulin (AREG) and ERBB3 in the integrated dataset.

**Fig. A6:**
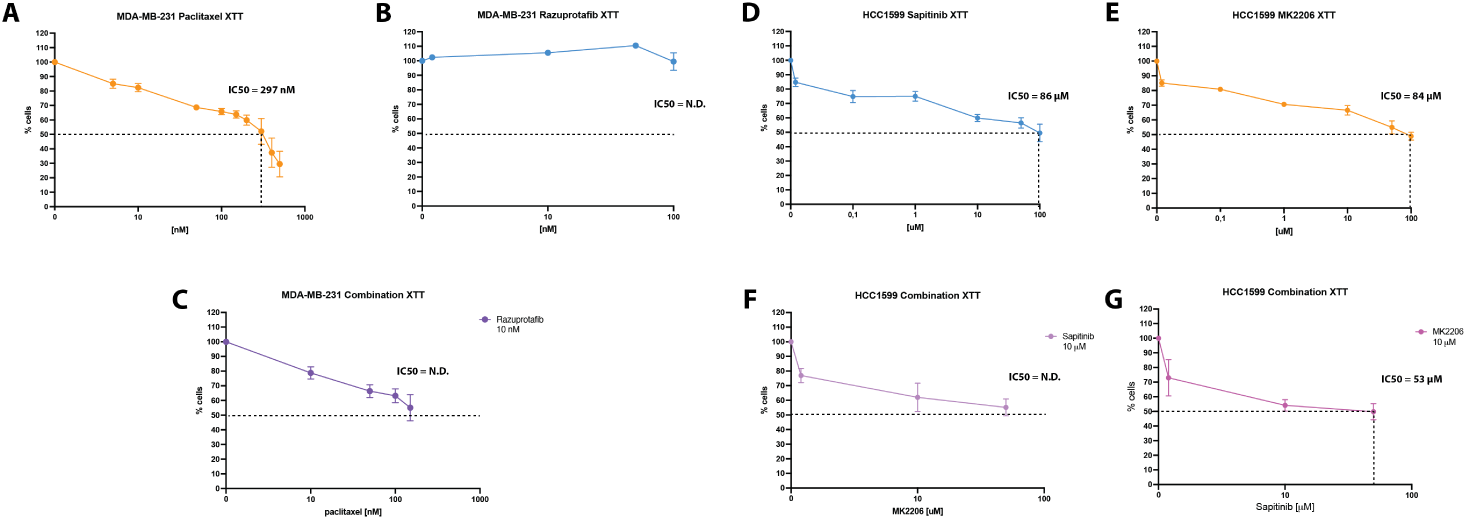
Dose response curves for MDA-MB-231 and HCC1599 in monolayer or suspension monocultures. **(A)** Dose response curve for paclitaxel treatment in MDA-MB-231 2D monocultures. **(B)** Dose response curve for razuprotafib treatment in MDA-MB-231 2D monocultures. **(C)** Dose response curve for combination treatment in MDA-MB-231 2D monocultures. **(D)** Dose response curve for sapitinib treatment in HCC1599 suspension monocultures. **(E)** Dose response curve for MK2206 treatment in HCC1599 suspension monocultures. **(F)** Dose response curve for combination treatment in HCC1599 suspension monocultures. **(G)** Dose response curve for combination treatment in HCC1599 suspension monocultures. N.D. = not determined

